# 3D Brain Vascular Niche Model Captures Invasive Behavior and Gene Signatures of Glioblastoma

**DOI:** 10.1101/2024.07.09.601756

**Authors:** Vivian K. Lee, Rut Tejero, Nathaniel Silvia, Anirudh Sattiraju, Aarthi Ramakrishnan, Li Shen, Alexandre Wojcinski, Santosh Kesari, Roland H. Friedel, Hongyan Zou, Guohao Dai

## Abstract

Glioblastoma (GBM) is a lethal brain cancer with no effective treatment; understanding how GBM cells respond to tumor microenvironment remains challenging as conventional cell cultures lack proper cytoarchitecture while *in vivo* animal models present complexity all at once. Developing a culture system to bridge the gap is thus crucial. Here, we employed a multicellular approach using human glia and vascular cells to optimize a 3-dimensional (3D) brain vascular niche model that enabled not only long-term culture of patient derived GBM cells but also recapitulation of key features of GBM heterogeneity, in particular invasion behavior and vascular association. Comparative transcriptomics of identical patient derived GBM cells in 3D and *in vivo* xenotransplants models revealed that glia-vascular contact induced genes concerning neural/glia development, synaptic regulation, as well as immune suppression. This gene signature displayed region specific enrichment in the leading edge and microvascular proliferation zones in human GBM and predicted poor prognosis. Gene variance analysis also uncovered histone demethylation and xylosyltransferase activity as main themes for gene adaption of GBM cells *in vivo*. Furthermore, our 3D model also demonstrated the capacity to provide a quiescence and a protective niche against chemotherapy. In summary, an advanced 3D brain vascular model can bridge the gap between 2D cultures and *in vivo* models in capturing key features of GBM heterogeneity and unveil previously unrecognized influence of glia-vascular contact for transcriptional adaption in GBM cells featuring neural/synaptic interaction and immunosuppression.

## INTRODUCTION

Glioblastoma (GBM) is the most aggressive and lethal form of brain cancer, with a median survival of only 15-18 months despite multimodal treatment ^1^. A major challenge for GBM research and therapy development has been the inability of traditional two-dimensional (2D) cell culture systems to recapitulate the complex tumor microenvironment (TME) and GBM heterogeneity observed *in vivo* ^2^. The lack of proper three-dimensional (3D) cytoarchitecture thus represents a major limitation to understand tumor biology and drug response ^3^.

Traditionally, GBM research has relied on animal models, such as patient derived xenografts (PDX) and genetically engineered mouse glioma models ^4^. While these *in vivo* models have provided valuable insights, they present the complexity of TME all at once, making it challenging to stepwise dissect the intricate interactions between tumor cells and stromal cells and with the surrounding neuronal and glial networks. The GBM microenvironment is highly complex and dynamic, characterized by unique extracellular matrix (ECM), dense but dysfunctional tumor vasculature, and abundant non-neoplastic stromal cell types such as glia and immune cells ^5–7^. These components affect tumor progression, invasion, immune suppression, and therapy resistance ^8–10^. Recreating the intricate TME in the laboratory setting faces major hurdles, as it requires the integration of multiple cell types, the establishment of appropriate cell-cell and cell-matrix interactions, and the incorporation of physiologically relevant mechanical and biochemical cues. Emulating these brain-specific characteristics is essential for teasing out the unique tumor biology and heterogeneity of GBM regarding invasion behavior, vascular association, transcriptional adaptations, proliferative *vs*. quiescent state, and therapy resistance. A better understanding many reveal novel molecular targets to disrupt the malignant GBM-neural network and counter immunosuppression.

To address the limitations of current GBM models, there is a growing need for biomimetic 3D culture systems with tissue-level complexity mimicking the brain microenvironment. Brain organoids and organ-on-a-chip platforms have emerged as promising tools, as they provide a more realistic representation of the tumor-host interactions to study angiogenesis, drug delivery, and therapy resistance ^11–13^. However, current brain organoids lack a functional vasculature, thus limiting their ability to fully mimic the complex vascular network of the brain ^14^. The organ-on-a-chip platforms have the limitations of inadequate ECM volume and constrained size for accurate reproduction of tumor growth dynamics, including the formation of hypoxic, necrotic zones, and the heterogeneity and invasiveness of tumor cells^15–17^.

In this study, we developed an improved 3D brain vascular model by incorporating human sourced brain-specific endothelial cells, pericytes, astrocytes and patient derived GBM cells. We conducted direct comparisons of tumor cell behavior, including invasion pattern and vascular association, as well as transcriptomic profiles across traditional 2D culture, various 3D models, and an orthotopic *in vivo* intracranial transplant model using the same patient derived GBM stem cells (GSC). Our results demonstrate that the advanced 3D brain vascular niche model can recapitulate key features of GBM heterogeneity and gene signatures concordant with invasion patterns and GBM survival. Together, these findings highlight the value of advanced 3D models as an effective bridge between 2D culture and *in vivo* transplants for GBM modeling and identification of new molecular targets to combat this devastating disease.

## RESULTS

### Optimization of 3D human brain vascular niche model using a multicellular approach

To build a biomimetic brain vascular niche model, we seeded in 3D fibrin matrix various combinations of human brain endothelial cells (bECs), human astrocytes (AC), brain pericytes (PC), and monitored vascular network formation (**Fig. 1a**). When co-cultured with astrocytes alone, bECs tended to form broad vacuoles with few interconnections; when seeded with brain pericytes, bECs formed interconnected vessels, but these vessels appeared thin and lacked hollow lumen formation. By contrast, the presence of both astrocytes and pericytes greatly enhanced vascular network connectivity, vessel width, hollow tube formation, and vascular area (**Fig. 1a**). This demonstrates the advantage of a multicellular approach for building a 3D brain vascular model.

**Figure 1.**
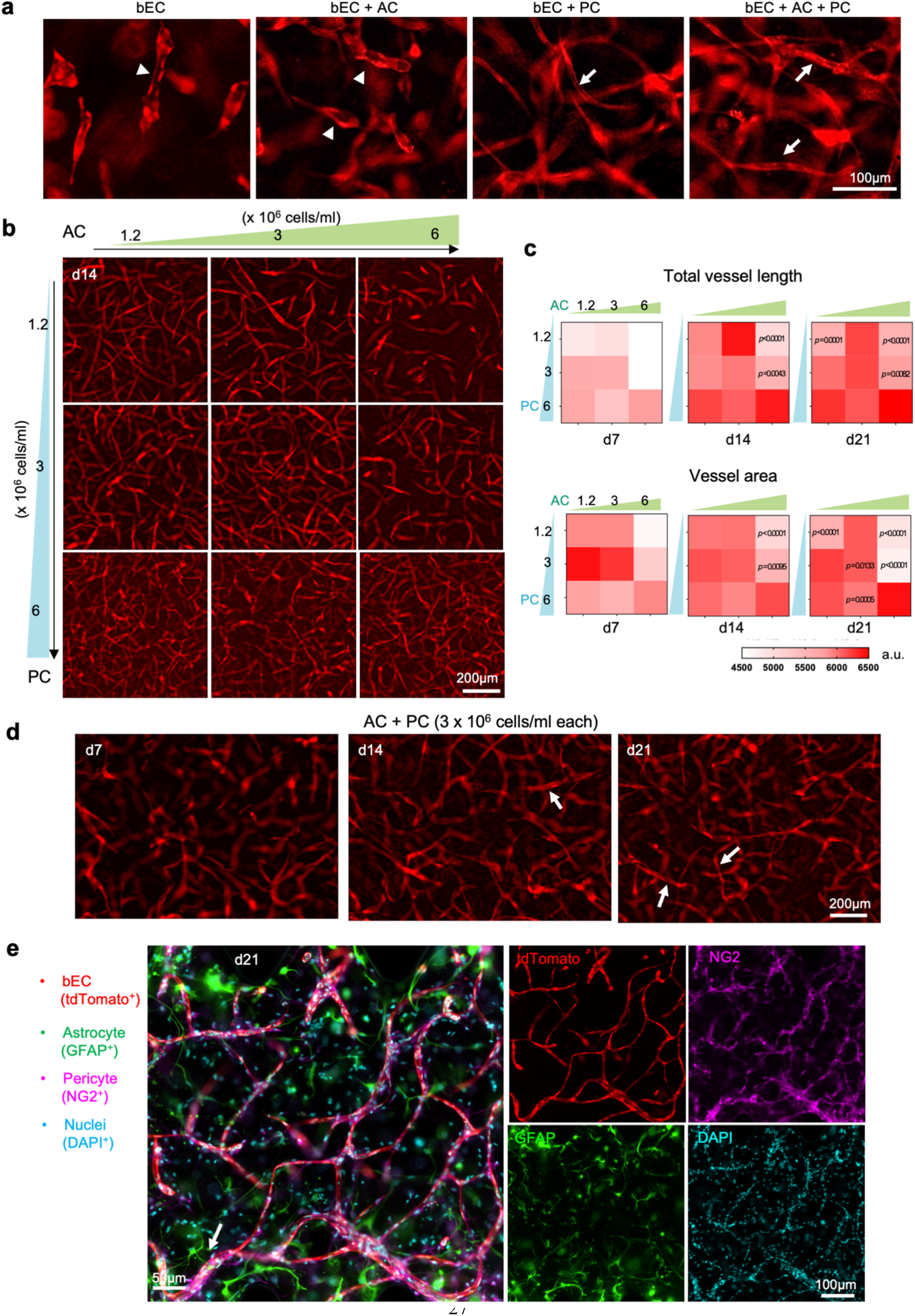
Optimization of a 3D brain vascular niche model. (a) Fluorescence live cell imaging of brain endothelial cells (bEC, tdTomato^+^), cultured alone in 3D gel or together with astrocytes (AC), pericytes (PC), or both at 14 day of culture. The bEC were seeded at a density of 6 x 10^6^ cells/ml and AC and PC at 3 x 10^6^ /ml. Note that bEC cultured alone or with AC only tended to form vacuoles that were not interconnected (arrowheads), while co-culture with PC, and more so with both AC and PC, led to the formation of interconnected vessels (arrows). (b) Representative images of vascular development of bEC (tdTomato^+^) in 3D gel when cultured with increasing seeding densities of AC and PC. (c) Quantifications of total vessel length and vascular area in cultures with different seeding densities of AC and PC. *p*-value calculated using one way ANOVA with Tukey’s post hoc test, compared to the AC6-PC6 condition (6 x 10^6^ cells/ml). n=4-12 independent cultures for each condition. (d) Time course of formation of interconnected vascular development of bECs (arrows) from culture day 7 to day 21, with AC and PC seeded at 3 x 10^6^ cells/ml each. (e) Co-immunofluorescence imaging of day 21 culture shows juxtaposition of PC (NG2^+^) with bEC (tdTomato^+^). Note AC (GFAP^+^) extending endfeet (arrow) towards vasculature. DAPI for nuclei staining.

We next fine-tuned the seeding densities of astrocytes and pericytes, testing different combinations of density for each cell type to further optimize vascular network formation (**Fig. 1b, c**). We found that high seeding density of astrocytes (6 x10^6^ cells/ml, referred to as AC6) led to considerable gel degradation and collapse of the 3D structure over time unless a high density of pericytes was also used. Increasing the seeding density of pericytes resulted in progressively longer vessel lengths, but a mid-range concentration (3 x10^6^ pericytes/ml, denoted as PC3) achieved a larger total vascular area (**Fig. 1b, c**). In total, five combinations of seeding densities – PC1.2-AC3, PC3-AC1.2, PC3-AC3, PC6-AC1.2, PC6-AC3) – resulted in high levels of both vessel length and vascular area at day 21 (**Fig. 1c, Fig. S1**), each with distinct vessel density and uniformity (**Fig. 1b**). For subsequent studies, we settled on the mid-range seeding densities of PC3-AC3 (3 x10^6^ cells/ml for both pericytes and astrocytes), which consistently yielded vasculature with abundant vessel branching and interconnected hollow tube structures after 14∼21 days of culture (**Fig. 1d**). Immunofluorescence staining further demonstrated perivascular localization of pericytes and astrocytes extending endfeet touching the blood vessels (**Fig. 1e**), reminiscent of the 3D cytoarchitecture of human brain microvascular network.

### 3D brain vascular niche model enables long-term culture of patient derived GBM stem cells with distinct invasion patterns

To model GBM invasion, we compared 2D and 3D culture systems for long-term sustainability and capability to capture distinct infiltrative behavior of GBM cells (**Fig. 2a**). For the 3D models, we further compared seeding GBM cells as dispersed cells or as spheroid aggregates, either in 3D fibrin hydrogel alone (3D G) or in the optimized 3D human brain vascular niche model containing also astrocytes, pericytes, and bECs (3D GAPE) (**Fig. 2b**). Two patient-derived GBM stem cell (GSC) lines were tested (SD2 and SD3), characterized as proneural and mesenchymal GBM transcriptional subtypes, respectively^18, 19^.

**Figure 2.**
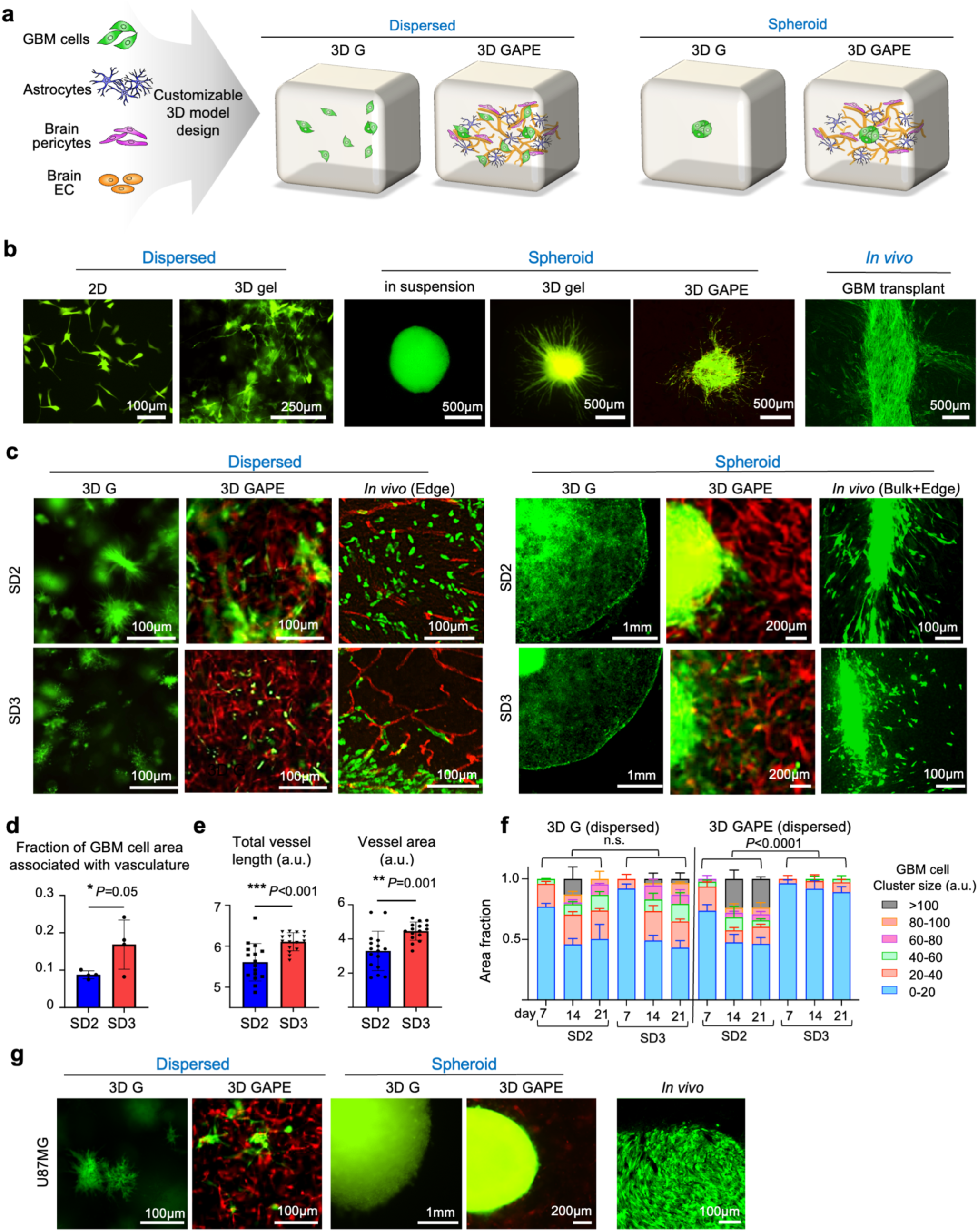
3D GAPE model recapitulates GBM invasion patterns as observed in vivo. (a) Schematics of 3D GBM model fabrication with GBM cells seeded either in dispersed or spheroid fashion, together with astrocytes, and brain-derived pericytes and endothelial cells (EC). Formation of a brain vascular network within the 3D structure with close interaction with GBM cells occurred over 3 weeks of culture. (b) Examples of patient derived GBM stem cells (expressing GFP) in various 2D or 3D cultures or in intracranial in vivo transplant in SCID mice. For 3D gel model, increased amount of matrix was used to sustain 3D gel model for over 2 weeks of culture. (c) Immunofluorescence images of patient derived GBM stem cells (GFP^+^) seeded in different 3D culture conditions in dispersed or spheroid fashion, as compared to in vivo intracranial transplants (human nuclear antigen^+^ in tumor edge, or GFP^+^ in bulk+edge). Both SD2 and SD3 exhibited similar growth patterns in the 3D G condition, but distinctive invasion patterns in the 3D GAPE model, with SD3 showing stronger vascular association, resembling *in vivo* invasion at tumor margin. Vasculature was visualized by tdTomato^+^ ECs in 3D GAPE model or by staining for PECAM1 for *in vivo* model. (d) Quantification shows a higher fraction of SD3 GSCs associated with vasculature in the dispersed 3D GAPE model (day 21) compared to SD2 GSC. n=4 independent cultures for each condition. Data represent mean ± SD. (e) Quantifications show higher total vessel length and vessel area for dispersed 3D GAPE model with SD3 GSCs compared to 3D GAPE with SD2 GSC (day 21). Unpaired two-tailed Student’s *t* test, n=16 independent cultures for each condition. Bar graphs represent mean ± SD. (f) Quantifications show differences in sizes of aggregates of SD2 and SD3 GSC in dispersed 3D GAPE model, but not in dispersed 3D G model. One way ANOVA with Tukey’s post hoc test. n=2 independent cultures for 3D G, n=4 independent cultures for 3D GAPE. Bar graphs represent mean ± SD. (g) Fluorescence images show growth pattern of human GBM cell line U87MG (GFP^+^) seeded as dispersed or spheroid cells in different 3D models or in in vivo intracranial transplant. Note that U87MG expanded as bulk mass without invasive pattern both in 3D and in in vivo models.

In the dispersed 3D G model without vasculature, both SD2 and SD3 GSCs formed small tumor clusters with limited individual cell migration or morphological differences between the two (**Fig. 2c**). However, when cultured in the dispersed 3D GAPE model, we detected morphological variations, with SD3 displaying a stronger vascular association than SD2 (**Fig. 2c, d**). This recapitulated *in vivo* migratory behaviors in intracranial transplants wherein both GSC lines displayed aggressive invasion but SD3 showed a preference for vascular association (**Fig. 2c**). Aside from differences in vascular association, SD3 also showed a propensity for individual cell invasion in the dispersed 3D GAPE model, in contrast to the collective migration favored by SD2 (**Fig. 2c, e**). Quantification of the size of GBM cell clusters confirmed a significant difference between SD2 and SD3 in collective vs. individual cell invasion in the 3D GAPE model, a phenotypic difference not captured in the 3D G model (**Fig. 2e**).

When cultured as GBM spheroids in the 3D G model, both SD2 and SD3 exhibited aggressive invasive migration from the spheroid edge, thus offering a clearer readout of invasiveness than the dispersed model with GBM cells scattered throughout the 3D cultures (**Fig. 2c**). However, by day 21, most of the gel structures of the spheroid 3D G model had been degraded by the GBM cells (**Fig. 2c**), and more volume of fibrin matrix was needed to maintain a 3-week culture. In contrast, in the spheroid 3D GAPE brain vascular niche model, matrix degradation was minimal and individual GBM cell invasion from the spheroid edge was clearly detected. Similar to the dispersed 3D GAPE model, in the spheroid 3D GAPE model we also observed a more robust vascular association of SD3 than SD2, again recapitulating the *in vivo* perivascular migratory behavior of SD3 (**Fig. 2c**). Another noticeable difference between the two GSC lines in the 3D GAPE model was an enhanced vessel development for SD3 as compared to SD2, measurable by both longer total vessel length and larger vascular area (**Fig. 2f**). Together, these data support the advantage of the 3D GAPE over the 3D G system to model GBM invasion, for both dispersed and spheroid models, with the capacity to reveal morphological differences between different patient GSC lines, including association with brain vasculature and collective *vs*. individual cell migration.

To further confirm the capability of our 3D GAPE model to capture *in vivo* GBM invasion behavior, we also compared GSC with U87MG cells, a widely used human GBM cell line that expands as a bulk mass tumor rather than by infiltrative growth ^20^. Indeed, mirroring the *in vivo* bulk growth mode in intracranial transplants, when seeded in 3D GAPE, U87MG cells expanded as a spheroid mass without infiltration along vasculature (**Fig. 2g**). Of note, in the 3D GAPE model, U87MG cells markedly suppressed vascular formation surrounding the GBM spheroid (**Fig. 2g**).

### GBM cells in 3D vascular model and in in vivo transplants share gene signatures featuring neural/glial interactions

Interactions with the TME can induce profound transcriptional changes in GBM cells. We therefore investigated next how GBM cells adjust gene expression in response to different culture conditions as compared to the *in vivo* transplant paradigm. To this end, we isolated SD2 and SD3 cells from 2D and 3D culture models and from *in vivo* intracranial transplants and conducted RNA-sequencing with three independent samples for each condition (**Fig. 3a, S2a**). For the 3D GAPE model, GBM cells were isolated from other cells by FACS for GFP fluorescence, and for *in vivo* transplants, GBM cells were separated from stromal cells by FACS using an antibody against human leukocyte antigen (HLA) (**Fig. S2b**). Quantitive qRT-PCR analysis for *CDH5* (VE-Cadherin), a EC marker, confirmed absence of its expression in all the GBM cell samples (**Fig. S2c**).

**Figure 3.**
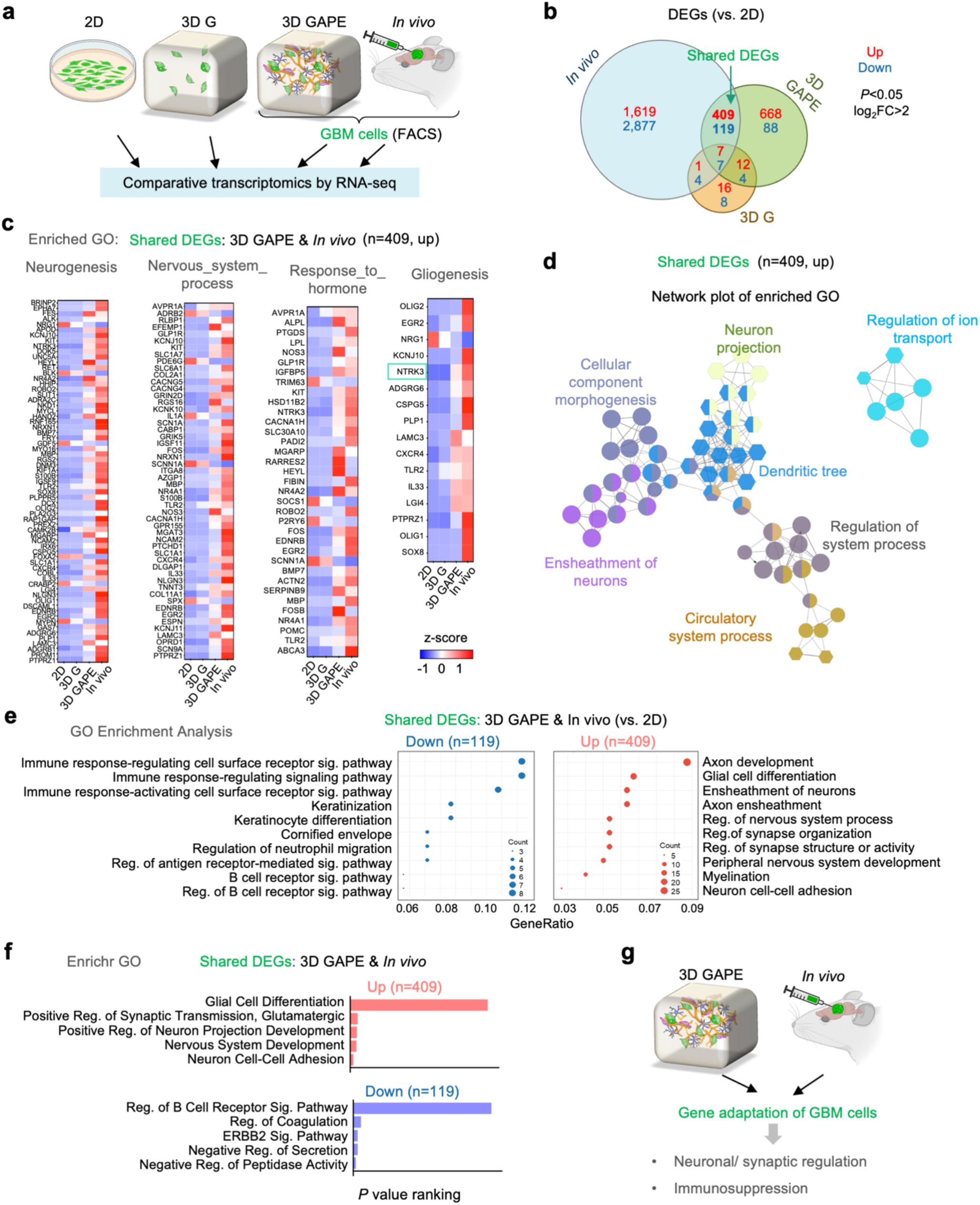
Comparative transcriptomics reveal shared differentially expressed genes in 3D GAPE and *in vivo* featuring neural/synaptic regulation and immunosuppression. (a) Schematic of RNA sequencing analyses of GBM cells grown in different 2D or 3D conditions or in intracranial transplants. GBM cells from 3D GAPE or *in vivo* transplants were isolated by FACS for GFP^+^ cells or HLA^+^ cells, respectively. (b) Venn diagram illustrating overlap of differentially expressed genes (DEGs, cutoff: *P*<0.05 and |log_2_fold change| >2) in GBM cells from different conditions relative to 2D. (c) Heatmap of expression of genes in top GOs enriched in upregulated DEGs shared by 3D GAPE and *in vivo* relative to 2D. (d) Network plot of enriched GO of shared upregulated DEGs of 3D GAPE and *in vivo* conditions relative to 2D. Circular nodes depict GO Biological Process, while hexagonal nodes indicate GO Cellular Component. (e) GO Enrichment Analysis of shared DEGs of 3D GAPE and *in vivo* relative to 2D, separated into up- and down-regulated DEGs. (f) Enrichr analysis of shared DEGs, separated into up and downregulated genes in 3D GAPE and *in vivo* relative to 2D. (g) Summary depiction of gene adaptation of GBM cells in 3D GAPE and *in vivo* featuring neural/synaptic regulation and immunosuppression.

We first identified differentially expressed genes (DEGs) of combined SD2 and SD3 GSCs in different 3D and *in vivo* models *vs*. 2D condition (**Fig. S3a**). This revealed that GBM cells *in vivo* harbored the highest number of DEGs relative to 2D, with more down-regulated than up-regulated genes (**Fig. 3b**). This was followed by 3D GAPE and 3D G as a distant third (**Fig. 3b**). Intersection analysis revealed far more common DEGs shared between *in vivo* and 3D GAPE than between *in vivo* and 3D G (**Fig. 3b**). Hence, the numbers of DEGs and a large overlap between 3D GAPE and *in vivo* support a better biomimetic environment conferred by 3D GAPE than the simpler 3D G model in recapitulating GBM transcriptomic adaptations.

Remarkably, the DEGs shared between 3D GAPE and *in vivo* relative to 2D were mainly associated with CNS functions (e.g., Neurogenesis, Nervous system process, Gliogenesis) and Response to hormone (**Fig. 3c**). Core network analysis of these common DEGs further highlighted Neuron projection, Dendritic tree, Ensheathment of neurons, Regulation of system process, and Regulation of ion transport as the main themes (**Fig. 3d**). The comparative transcriptomic data thus illustrate an instructive role of the brain glia-vascular unit (containing astrocytes, brain pericytes, and brain ECs) in inducing gene programs in GBM cells for neural interactions and synaptic regulation, despite the absence of neurons in the 3D GAPE model.

We next analyzed the up- and down-regulated common DEGs separately. Enrichr gene ontology (GO) enrichment analysis revealed that, as compared to 2D condition, the environments in 3D GAPE or *in vivo* induced genes related to neuronal development (i.e., axon development, neuron cell-cell adhesion, peripheral nervous system development), glial cell differentiation and myelination (i.e., ensheathment of neurons, axon ensheathment, myelination), and regulation of synapse structure or activity (**Fig. 3e, S3b**). In echo, Enrichr analyses further showed that the shared up-regulated DEGs between 3D GAPE and *in vivo* were highly enriched for pathways concerning neuronal interactions and nervous system development (**Fig. 3f**). By contrast, the shared down-regulated DEGs largely concerned immune signaling (*e.g*., regulation of B cell receptor signaling pathway, regulation of antigen receptor-mediated signaling pathway, regulation of neutrophil migration), in accordance with the well-known immunosuppressive status of GBM (**Fig. 3e, f**). Together, these initial comparative transcriptomics data showcase the biomimetic nature of our 3D GAPE microenvironment in inducing gene programs featuring neural/glial interaction as well as immunosuppression (**Fig. 3g**).

### Shared gene signatures of GBM cells in 3D vascular culture and *in vivo* predict poor outcome for GBM patients

We further analyzed the upregulated DEGs shared by GBM cells in 3D GAPE and *in vivo*, which were associated with 5 top enriched GO categories (ranked by gene ratio) related to axon development, axon ensheathment, glial cell differentiation, ensheathment of neurons, and regulation of nervous system process (**Fig. 4a, b**). For instance, the shared DEGs included *UNC5A, MGARP, MT3, KIAA1755, NCAM2, CSPG5, EGR2, NR4A2, S100B, DOBL, ARK2C,* genes that are linked to axon development. The shared DEGs also included *NTRK3, LAMC3, SOX8, ERBB3, OLIG1, LGI4, TLR2, OLIG2, KCNJ10, CSCR4, ADGRG6, CLDN11, MPZ, GAL3ST1,* genes related to ensheathment of neurons, glial cell differentiation and/or axon ensheathment. Genes linked to regulation of nervous system process such as *CACNG4, GRIN2D, AVPR1A, NOS3, IGSF11, OPRD1, DLGAP1, CACNG5, EDNRB* were also included in the shared DEGs. *CSPG5, MT3, PLP1, PTPRZ1, MBP, EGR2, IL33, LGI4, TLR2, OLIG2, CXCR4, CLDN11, MPZ, GAL3ST1, NLGN3, NRXN1* were genes concurrently associated with more than two categories mentioned above (**Fig. 4a, b**). Altogether, GBM cells adapted gene expression for neural/glial interactions in response to the biomimetic glia-vascular niche environment provided by our 3D GAPE model.

**Figure 4.**
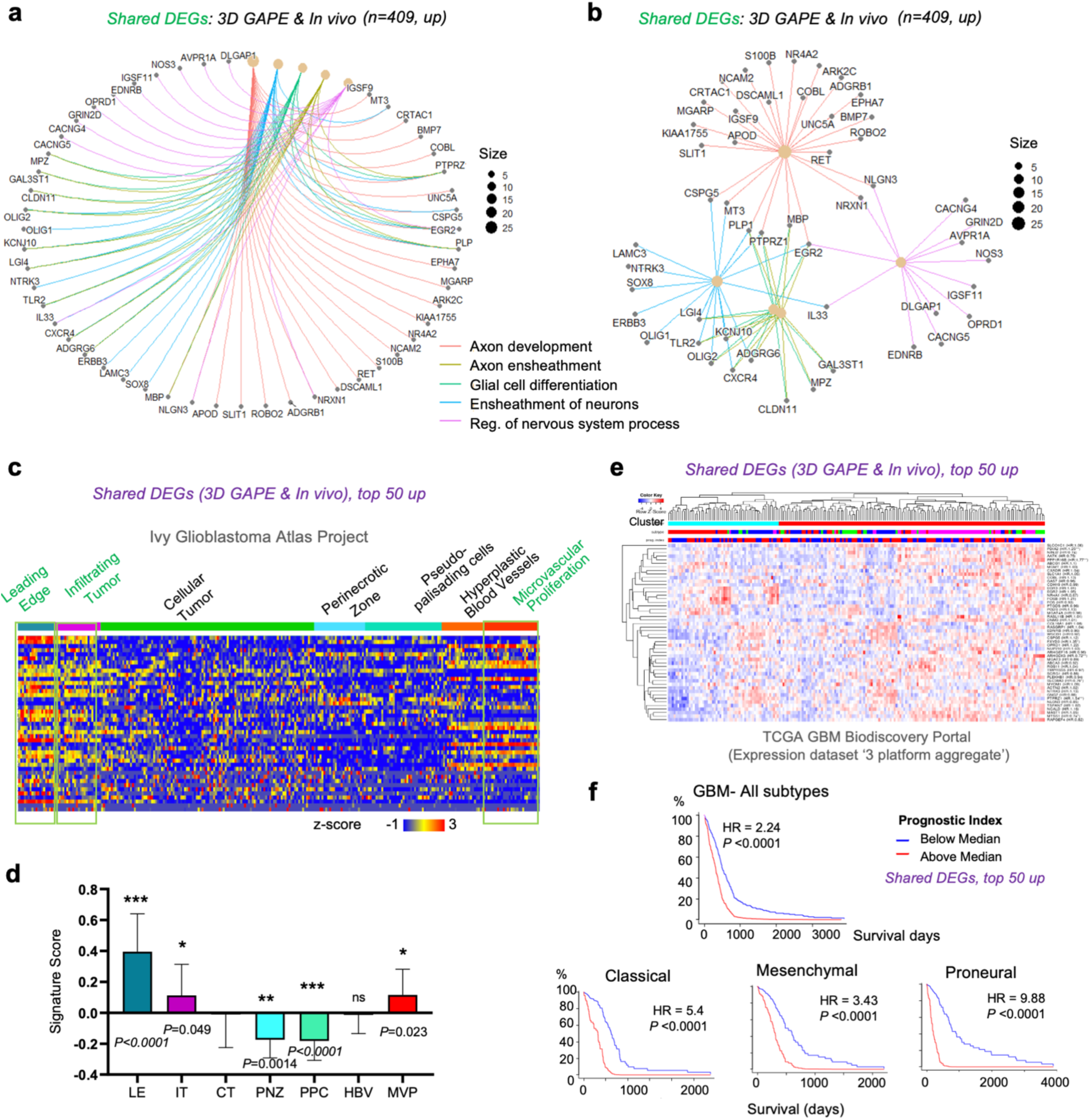
Shared GBM gene signatures of 3D GAPE and *in vivo* are expressed at distinct zones of human GBM and predict poor survival. (a, b) Gene-concept network depiction of 5 top enriched GOs (ranked by *p-*value) of the upregulated DEGs in 3D GAPE and *in vivo* relative to 2D (n=409). The layout displays the relationships between significantly enriched GO terms (colored nodes with predicted biological functions) and their associated genes. (c, d) Heatmap and quantification of the expression of 43 (of 50) top upregulated DEGs shared by 3D GAPE and *in vivo* relative to 2D in different zones of human GBM patient samples (Ivy GAP database). Note significantly higher expression of the shared DEGs in leading edge and infiltrating tumor zones, as well as microvascular proliferation zone. Mean expression scores for the 43 top shared DEGs across GBM tumor zones, normalized to the Cellular tumor (CT) zone. One-way ANOVA, followed by Tukey’s post hoc test. n=270 specimens (n=19-111 for each zone) in the Ivy GAP database. Data represent mean ± SD. (e) Top 26 (of 50) shared upregulated DEGs of 3D GAPE and *in vivo* were applied for cluster analysis of the TCGA GBM Biodiscovery portal. Note that only 26 of the top 50 genes were represented in Biodiscovery database. (f) Prognostic index based on expression of 26 (of 50) upregulated shared DEGs from 3D GAPE and *in vivo* in human GBM patients, shown for GBM in total and for individual transcriptional subtypes.

To assess the clinical significance of the shared DEG signature common to 3D GAPE and *in vivo*, we selected the top 50 upregulated genes ranked by *P* values and examined their expression pattern in human GBM by surveying the Ivy Glioblastoma Atlas Project, which contains region-specific gene expression data of human glioblastoma ^21^ (**Fig. 4c**). Strikingly, we found that the top 50 shared upregulated DEGs were significantly enriched in the tumor zones ‘leading edge’, ‘infiltrating tumor’, and ‘microvascular proliferation’, as compared to regions of GBM interior (‘cellular tumor’, ‘perinecrotic zone’, ‘pseudopalisading cells’) (**Fig. 4c, d**).

We next surveyed the TCGA GBM Biodiscovery Portal ^22^ for patient survival data, and found that the shared gene signature of 3D GAPE and *in vivo* was associated with poor prognosis for GBM patients, which was applicable to all three transcriptional GBM subtypes (**Fig. 4e, f**). These data underscore the clinical importance of the gene programs induced by glia-vascular contact to support neural/glial interactions and malignant potency of GBM.

### Variable genes across GBM models involve histone demethylation in response to 3D glia-vascular culture and *in vivo* environment

We next studied the variance of gene signatures of GBM cells in different conditions. Principal component analysis (PCA) revealed a clear distinction between SD2 and SD3 samples, consistent with the marked intertumoral heterogeneity of GBM (**Fig. 5a**). For both SD2 and SD3, the *in vivo* samples were distinctly separated from all three *in vitro* conditions, but distance to *in vivo* condition was closer for samples from 3D GAPE than from 2D and 3D G conditions (**Fig. 5b**).

**Figure 5.**
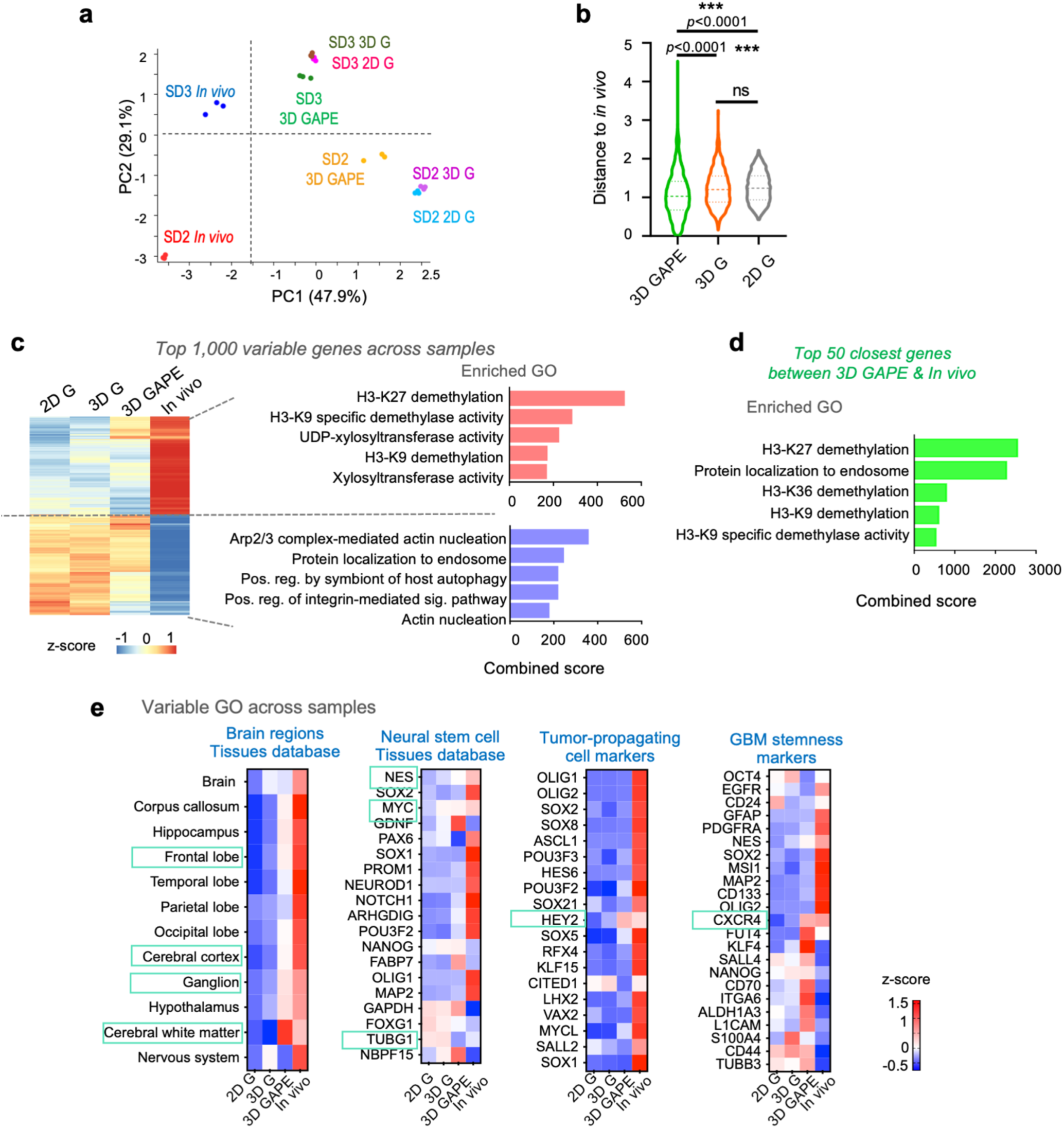
Variable genes highlight histone demethylation for epigenetic adaptation as a main theme shared by GBM cells in 3D brain vascular niche model and *in vivo*. (a) Principal component analysis (PCA) of RNA-seq samples show separation of *in vivo* samples from *in vitro* samples for both SD2 and SD3, but 3D GAPE samples being closest to *in vivo* samples on PC1 axis than other culture conditions. (b) Spearman correlation distances of cluster gene expression profiles between different in vitro conditions and *in vivo* samples (SD2 and SD3 combined), based on the top 1,000 most variable genes. Distances were calculated using hierarchical clustering. Violin plots show median, quartiles, and minimum and maximum values. (c) Left, unsupervised clustering of expression of top 1,000 most variable genes across SD2 and SD3 combined samples in different conditions. Right, enriched GOs of top 1,000 variable genes, separated into up and downregulated genes. (d) Gene ontology enrichment analysis of top 50 most closely correlated genes of GBM cells in 3D GAPE and *in vivo* conditions highlight epigenetic adaptations by histone demethylation as the main biological theme. (e) Top variable GO ontology terms across samples, based on geneset databases of brain regions and neural stem cells, as well as genesets associated with tumor propagation and GBM stemness.

The transcriptional distinction between *in vivo* and *in vitro* conditions was further illustrated by a heatmap of relative expression of the top 1,000 variable genes across samples (**Fig. 5c**). Notably, as compared to 2D or 3D G, the gene profile of 3D GAPE GBM cells appeared more similar to *in vivo*. Intriguingly, among the top variable genes, upregulated genes prominently concerned histone demethylation (for both H3-K27 and H3-K9) (**Fig. 5c**). Another top enriched GO term for the top upregulated variable genes was xylosyltranferase activity (**Fig. 5c**), which is linked to post-translational modification of proteoglycans such as CSPG with a major role in modulation of tumor biology ^23^. The top downregulated variable genes *in vivo* concerned cytoskeletal dynamics (*e.g*., Arp2/3 complex-mediated actin nucleation, integrin-mediated signaling pathway, actin nucleation), autophagy and endosome processes (**Fig. 5c**).

As another sign of a biomimetic microenvironment provided by our vascular niche model for GBM cells, the top 50 closest genes between 3D GAPE and *in vivo* paradigm were enriched for GO categories of histone demethylation and protein localization to endosome (**Fig. 5d**). We next further analyzed top variable GO terms across conditions, which revealed that the *in vivo* TME strongly induced genes linked to the nervous system (Brain regions Tissues database), in particular, frontal lobe, cerebral cortex, ganglion, and notably cerebral white matter, with 3D GAPE environment again most closely following this trend as compared to 2D or 3D G (**Fig. 5e**). The *in vivo* condition also strongly induced genes related to neural stem cell biology, tumor-propagation, and GBM stemness, including *SOX2*, *OLIG2*, *CD133*, *GFAP*, *MAP2, NESTIN, FUT4, KLF4, LICAM, CD44*, as well as tyrosine kinase receptors (*EGFR*, *PDGFRA*), all of which were not induced in 3D GAPE condition, highlighting a remaining gap between *in vivo* and *in vitro* paradigms (**Fig. 5e**). This also signifies a potential important impact of neuronal interactions on gene adaption for GBM stemness. Despite the absence of neurons in the 3D GAPE model, several neural stem cell markers such as *NES* and *MYC,* as well *TUBG1* (encoding tubulin gamma 1, involved in neuroprogenitor migration ^24^) were similarly induced in 3D GAPE and in *in vivo*. Likewise, as in *in vivo,* GBM cells in 3D GAPE also induced *HEY2*, a transcription factor of the Notch pathway, and chemokine receptor *CXCR4*, both linked to malignant potency of GBM ^25, 26^ (**Fig. 5e**).

### The 3D vascular model maintains a quiescent niche with protective role against chemotherapy

The vascular niche has been suggested as a key regulator of glioblastoma stem cells, with potential impact on GBM stem cell quiescence ^27, 28^. To assess the capacity of our model to maintain a quiescent niche for GBM cells, we leveraged a doxycycline (Dox) inducible histone 2B (H2B)-GFP quiescence reporter that we had previously developed ^19^, with quiescent or slow dividing cells retaining nuclear GFP signals while fast dividing cells lose the H2B-GFP signals over time (**Fig. 6a**). We induced H2B-GFP expression in GSC by Dox pulse from 7 days prior to seeding until 3 days after, and GFP signals were tracked during the following -Dox chase period until day 23 (**Fig. 6b**). While the initial GFP label retention patterns appeared similar across culture conditions at day 7, by day 15 and 21 days, the GFP signals largely disappeared in 2D G and 3D G condition; by contrast, we detected a significant amount of H2B-GFP^high^ signal in the 3D GAPE condition (**Fig. 6c, d**). Notably, quiescent GBM cells appeared to cluster together in perivascular niches (**Fig. 6e**).

**Figure 6.**
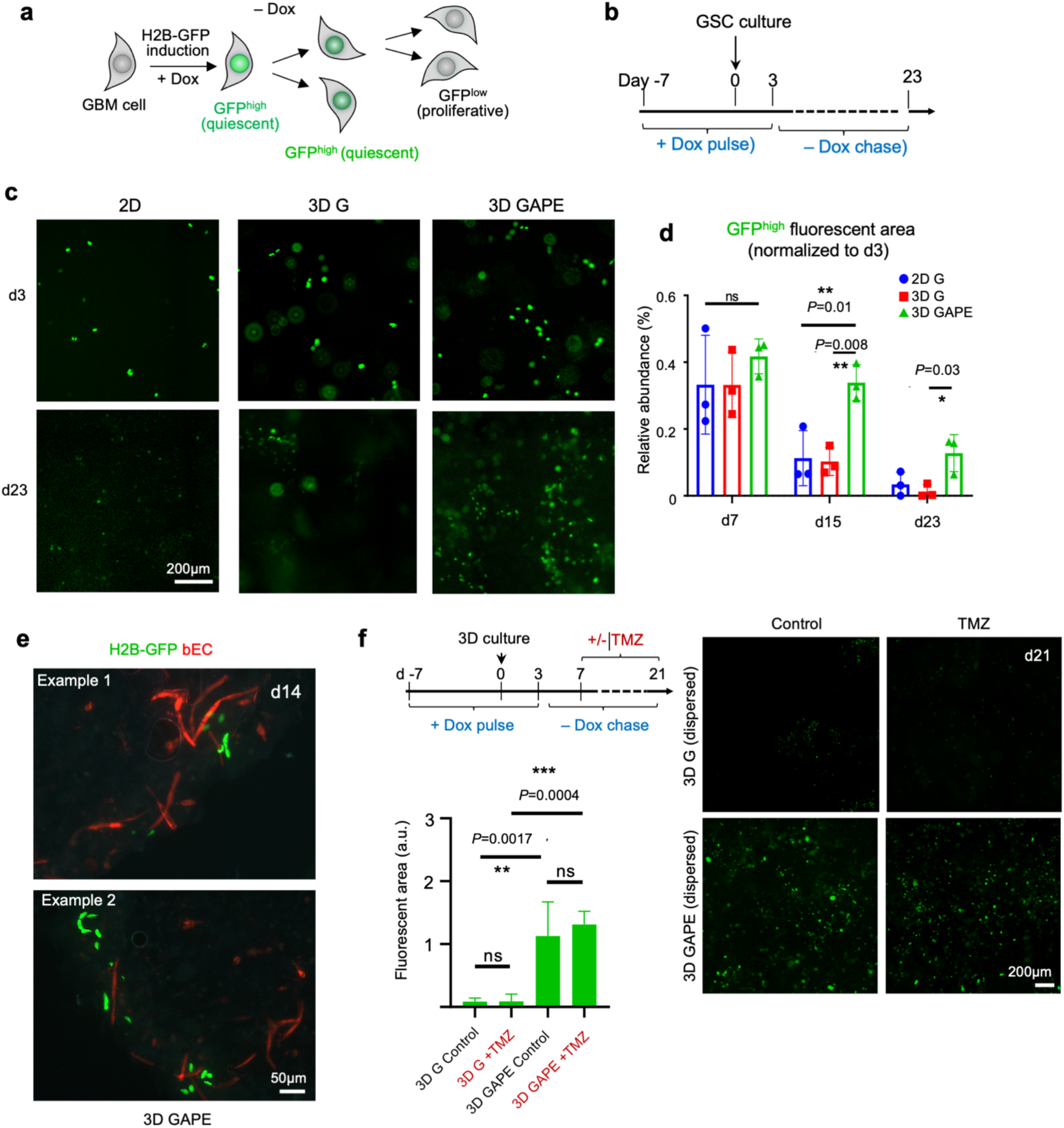
The 3D brain vascular model supports GBM quiescence niche. (a) Schematic of doxycycline (Dox)-induced expression of histone 2B(H2B)-GFP and progressive dilution of H2B-GFP label in proliferative cells during -Dox chase phase, while quiescent cells retain H2B-GFP label. (b) Experimental timeline of H2B-GFP induction (+Dox) and subsequent -Dox chase of GBM cell culture. (c) Fluorescence images of GBM cells in different culture conditions at day 3 and day 23. Note that day 3 was the last day of Dox pulse and day 23 was after 21 day of -Dox chase. (d) Quantification of GFP ^high^ fluorescent areas across culture conditions and culture periods. Note that GBM cells in 3D GAPE maintained a significantly higher amount GFP^high^ quiescent cells than other culture conditions. One-way ANOVA, followed by Tukey’s post hoc test. n=3 independent cultures for each condition. Bar graphs represent mean ± SD. (e) Two representative IF images show aggregation of H2B-GFP^high^ quiescent GBM cells in close proximity to vasculature at day 14. (f) Top left, timeline of H2B-GFP induction (+Dox) and treatment with chemodrug temozolomide (+/- TMZ) of 3D cultures from day 7 to day 21. Fluorescent images and quantification show significantly more H2B-GFP label-retaining GBM cells in 3D GAPE than in 3D G cultures at day 21, which survived even after TMZ treatment. One-way ANOVA, followed by Tukey’s post hoc test. n= 4 independent cultures for each condition. Bar graphs represent mean ± SD.

Quiescent GBM cells carry higher resistance to chemotherapy, which targets mostly proliferative cells^29^. For drug resistance studies, we administered from day 7 to day 21 during the Dox chase period temozolomide (TMZ), an alkylating chemo drug used as standard care for GBM patients (**Fig. 6f**). We observed that 3D GAPE maintained a similarly high number of H2B-GFP^high^ quiescent cells after two-week TMZ treatment as in vehicle-treated condition, demonstrating the chemodrug resistance of quiescent GBM cells (**Fig. 6f**). Hence the 3D GAPE microenvironment maintains a quiescent and also protective niche to resist chemotherapy.

## DISCUSSION

In this study, we optimized a biomimetic 3D brain vascular niche model for studying GBM, a highly aggressive brain cancer. The model incorporates key components of glia-vascular units with human-sourced brain mural cells and endothelial cells. Direct comparison of patient derived GSC lines across 2D and 3D models with the *in vivo* intracranial transplant paradigm demonstrated capability of our model to capture key features of GBM heterogeneity such as invasion behavior and vascular association, gene reprogramming, as well as capacity to maintain a quiescent niche for GBM cells and protection against chemotherapy.

Creating a reproducible and sizable 3D environment for GBM modeling faces major challenges, particularly in maintaining long term matrix integrity. Our studies reveal that traditional 3D GBM spheroid cultures degrade the matrix rapidly, necessitating a large volume of matrix to sustain the culture^30^. By reducing the seeding density of GBM cells and employing a dispersed model, we mitigated matrix degradation issues. By contrast, the 3D GAPE model allowed long term culture of patient derived GSCs without significant matrix degradation, thus providing a stable and biomimetic microenvironment for long-term study of tumor cell behavior, gene adaptation, and drug response. For instance, SD3 displayed a predilection for vascular association than SD2, detectable in both 3D GAPE and in PDX *in vivo* models, but not in 3D G condition.

We compared three different human GBM cells: patient-derived SD2 and SD3 cells with stemness characteristics, representing proneural and mesenchymal GBM subtypes, respectively ^19^, and the U87MG cell line. This provided compelling results on the capability of our 3D brain vascular niche model to recapitulate key features of GBM intertumoral heterogeneity, including invasion behavior (collective migration *vs*. single cell infiltration *vs*. no invasion), vascular association, and transcriptional changes in response to TME. The ability to directly compare our 3D model with both 2D culture and *in vivo* PDX model using identical GBM cells is particularly valuable, as it showcased the strengths and limitations of each system.

Remarkably, comparative transcriptomics across culture conditions versus *in vivo* unveiled a previously unappreciated influence of brain glia-vascular unit (consisting of astrocytes, brain pericytes and brain EC) in gene adaptation of GBM cells that featured neurogenesis, glial differentiation, and nervous system processes. Intriguingly, despite the absence of neurons or immune cells in the 3D vascular model (inclusion of neuron and immune cells will be a future research direction), the GBM gene signature shared by 3D GAPE and *in vivo* conditions highlighted synaptic regulation and immunosuppression. This insight may provide a new therapeutic angle by way of targeting tumor vasculature to disrupt the malignant connection between GBM cells and neuronal network. This is particularly relevant in light of recent findings on a promoting role of neuronal activity for GBM proliferation and progression ^31^. It may also provide an alternative strategy to counter the immunosuppressive state that is notorious for GBM, based on our findings that contact of GBM cells with glia-vascular units triggered downregulation of immune signaling genes. Indeed, the gene signatures shared between 3D GAPE and *in vivo* showed a regional enrichment in specific zones of human GBMs, in particular leading edge, infiltrating tumor, and microvascular proliferation zones, and predicted poor survival for GBM patients, thus further supporting clinical relevance of our model for understanding malignant potency of GBM.

Previously, to mimic the stiffness of brain microenvironment, various biomaterial hydrogels were developed to culture GBM cells ^32–35^. However, our results showed that hydrogel alone (3D G) falls short of creating a representative tumor microenvironment, as it could not capture *in vivo* GBM gene signature adaptations and several other key features of GBM. Our results underscore the importance of a cellular microenvironment in dictating GBM phenotypes. To address the cellular components, earlier studies predominantly relied on human umbilical vein endothelial cells (HUVECs) and endothelial cells from other sources ^36–38^. Recently, brain relevant vascular cells have been introduced to construct GBM models ^33, 39–42^. However, most prior studies focused on *in vitro* characterization but did not compare to gene signatures induced in the *in vivo* condition. We employed here brain-specific cells to construct a 3D vascular niche model, as brain ECs display unique functional features ^43, 44^. The use of all brain-sourced human cells may be crucial to induce gene reprogramming concerning nervous system processes. Our transcriptomics studies comparing GBM cells across different *in vitro* and *in vivo* conditions is powerful in unveiling previously unrecognized gene pathways that prominently featured histone demethylation and xylosyltransferase activity. Future studies are needed to understand the functional significant of the epigenetic changes and post-translational modification of proteoglycans such as CSPG in GBM cells in response to TME.

Finally, utilizing GSCs engineered with a quiescence reporter, we demonstrated the capability of the 3D GAPE culture model to maintain a quiescent and a protective niche against chemotherapy, which was not observed in 2D G and 3D G models. TMZ treatment in 3D GAPE resulted in an almost intact GFP^high^ area, suggesting that the quiescent cells may be the predominant population that display chemo resistance. Further isolating and characterizing this cell population may reveal important molecular mechanism of dormancy and therapeutic resistance. Our 3D GAPE *in vitro* model not only offers a platform for investigating GBM pathophysiology but can also serve as a customizable tool for drug discovery and studying vascular interactions, such as blood-brain barrier (BBB) function ^45^. Future directions would also incorporate neurons and immune cells in our models, which will be valuable to understand the network between malignant GBM cells and neuronal synaptic connectivity and immunosuppression mechanisms.

In summary, our study highlights the potential of an advanced 3D brain vascular niche model to serve as a bridge between traditional culture systems and animal-based *in vivo* approaches in capturing GBM heterogeneity and providing a versatile and a more physiologically platform to study GBM biology and advance drug discovery.

## MATERIALS AND METHODS

### Cell Culture

Fibrinogen and thrombin (Millipore Sigma) were dissolved in culture media or Dulbecco’s phosphate-buffered saline (DPBS) for 3D matrix fabrication. Temozolomide and doxycycline hyclate were purchased from Millipore Sigma. Human brain microvascular endothelial cells (Sciencell) were transduced with a lentivirus to express tdTomato fluorescent protein (Vectorbuilder) and were cultured in EGM-2 media (Promocell or Lonza). Human astrocytes (Sciencell) and human brain vascular pericytes (Sciencell) were expanded and cultured in astrocyte medium (Sciencell) and pericyte medium (Sciencell), respectively.

Patient-derived glioblastoma stem cell lines SD2 and SD3 ^19^ were transduced with GFP-expressing lentivirus or were genetically engineered with a doxycycline-inducible H2B-GFP reporter ^19^ and cultured in laminin-coated tissue culture flasks in complete NeuroCult NS-A proliferation medium for human cells (Stemcell Technologies). U87MG cells (ATCC) were transduced with a GFP expressing lentivirus and cultured in Dulbecco’s modified Eagle’s medium (DMEM; Gibco) supplemented with 10% fetal bovine serum and penicillin/streptomycin. To create GBM cell spheroids, 1,000 to 5,000 GBM cells were plated into a Corning spheroid microplate and cultured for 5 to 10 days until spheroids reached the desired diameter of >400 µm.

### Modelling of brain vascular niche in 3D culture

3D fibrin matrices containing varying numbers of brain endothelial cells (bEC), astrocytes (AC), and pericytes (PC) were prepared by the following method. Fibrinogen solution containing cells was prepared at a concentration of 20 mg/ml in warm EGM-2 media. Thrombin solution was prepared in DPBS at a concentration of 6 u/ml. The fibrinogen solution containing cells (20 µl) and thrombin solution (20 µl) were mixed through gentle pipetting and deposited on culture plates. The final seeding density of bEC-tdTomato was 6 x10^6^ cells/ml for all samples. The final seeding density of AC and PC were 1.2, 3, or 6 x10^6^ cells/ml. The 3D samples were cultured in EGM-2 media for 21 days. Live imaging was performed every 3-4 days using Nikon Eclipse Ti2 microscope to observe vessel growth over time. Quantification of vessel area and total vessel length were performed using ImageJ software.

To establish a brain vascular niche incorporating GBM cells in dispersion, GBM cells were mixed into fibrinogen solution containing bEC, AC, and PC at a density of 25,000 cells/ml. This mixture was then combined with the thrombin solution to facilitate crosslinking. For the GBM spheroid model, a GBM spheroid was placed into a culture plate well using a wide-bore pipette tip, followed by the removal of excess liquid. The fibrinogen solution containing bEC, AC, and PC was mixed with the thrombin solution, and the resulting blend was injected beneath the GBM spheroid to elevate it slightly. As the fibrin crosslinked, the GBM spheroid sank and became positioned at the center of the 3D hydrogel structure. All 3D GBM-vascular niche samples were cultured in EGM-2 media for up to 21 days, with media changes occurring every 2-3 days. Live imaging was performed every 3-4 days using Nikon Eclipse Ti2 microscope to observe GBM cell growth/invasion within the vascular niche.

### In vivo GBM intracranial transplant model

All animal procedures were approved by the Institutional Animal Care Use Committee (IACUC) of Icahn School of Medicine at Mount Sinai. Adult 8-week-old ICR-SCID mice (Taconic) were anesthetized with isoflurane (5% initial, then 1% continuous), and 2-3 x10^5^ GBM cells (SD2 or SD3 GSCs) were injected into each the right and the left striatum of recipient mice with a microsyringe (Hamilton) attached to a stereotactic instrument (Stoelting) (coordinates: 2 mm right/left and 0.5 mm anterior of Bregma, 2 mm deep). Mice bearing SD2 transplant were euthanized after 18-19 weeks, and mice bearing SD3 transplants after 5-7 weeks, and brains were removed for further analysis.

### FACS sorting

For 3D G and 3D GAPE in vitro cultures, GFP-expressing GBM cells were isolated by degrading the 3D matrix by a brief trypsin-EDTA treatment and dissociation in accutase, followed by sorting for GFP fluorescence using a BD FACSAria device. After extracting total RNA from sorted cells, expression of *CDH5* (VE-cadherin) was determined via PCR to confirm absence of ECs in the sorted population.

Human GSCs transplanted into mouse brain were isolated by dissociation of forebrain tissues with the Neural Tissue Dissociation kit (Papain; Milltenyi 130-092-628) and resuspended in FACS buffer (Hibernate-E low fluorescence (BrainBits) with 0.2% BSA and 20 μg/ml DNase I (Worthington)) and passed through a 40 µm mesh filter into round-bottom tubes (Falcon). To separate human GBM cells from mouse host cells, cell suspension was incubated with the human-specific anti-human leukocyte (HLA) antibody (abcam #ab70328), and HLA^+^ cells were collected with a FACSAria IIu device (BD Biosciences) for subsequent RNA-seq (**Fig. S2b**).

### RNA-seq

Total RNA was extracted using RNeasy mini kit (Qiagen) and cDNA libraries for sequencing were prepared using NEBNext Ultra II Directional RNA Library Prep Kit for Illumina (NEB #E7760; ∼50-150 ng RNA input per replicate sample). RNA-seq was performed at John Wayne Cancer Institute using Illumina HiSeq2500 (1 x 75bp, 30 x10^6^ reads per sample).

Raw sequence reads from samples were mapped to the human genome (hg38) using HISAT2 ^46^. Counts of reads mapping to genes were obtained using HTSeq-count software against Ensembl v90 annotation^47^. Differential gene expression analysis was performed with the DESeq2 package ^48^.

Gene set enrichment analysis (GSEA) was performed using a pre-ranked list of gene expression fold change (FDR-corrected *P*<0.05) (http://software.broadinstitute.org/gsea/index.jsp). Network plots were generated using Cytoscape (https://cytoscape.org). For the analysis of genes correlated with TCGA GBM data, the top 50 upregulated genes (ranked by *p-*value) were analyzed using the TCGA GBM Biodiscovery Portal (glioma-biodp.nci.nih.gov).

### GSC Dormancy and Drug Treatment Resistance assays

SD2 and SD3 H2B-GFP GSCs cells were expanded in the presence of 1 µg/ml doxycycline for 14 days until for full expression of nuclear H2B-GFP, then harvested for 3D sample fabrication. GSC cells were mixed in a fibrinogen solution containing cells at 25,000 cells/ml. The fibrinogen solution was mixed with thrombin solution in 1:1 ratio, then deposited to culture plate before the crosslinking was completed. The fibrin crosslinking was achieved by mixing fibrinogen and thrombin solutions. Each sample was given EGM-2 with 1 µg/ml doxycycline for Days 0-3. Doxycycline was removed for Days 4-21 to allow for dilution of H2B-GFP signal by cell division.

For the treatment resistance assay, control and treatment group samples were cultured in EGM-2 media with doxycycline until Day 3, and afterwards without doxycycline. At Day 7, EGM-2 media was supplemented with 1.25 mM temozolomide chemotherapeutic in treatment groups. The temozolomide treatment was continued through Day 21. Live imaging was performed every 2-3 days using Nikon ECLIPSE Ti2 microscope. Endpoint samples were stained using viability indicator ethidium homodimer-1 (Invitrogen) and nuclear stain Hoechst 33342 (Invitrogen). The quantifications of GFP signal and of dead cells were performed using ImageJ software. One way ANOVA was performed, followed by Turkey’s *t*-test, for statistical analysis.

### Immunofluorescent staining

Immunofluorescent staining techniques were used to fluorescently label specific proteins at the conclusion of cell culture. Briefly, all samples were washed three times with PBS and fixed with 4% paraformaldehyde (Alfa Aesa) for 1 hour at room temperature. Next, samples were washed three times with PBS and incubated with a blocking/permeabilizing (B/P) solution comprised of 10% normal goat serum (MP Biomedicals), 0.2% Triton X-100 (Fisher Scientific), and 0.1M glycine (Fisher Scientific) in PBS overnight at 4°C. Samples were then incubated with primary antibodies in B/P solution overnight at 4°C. PCs and ACs were stained with neural/glial antigen 2 (NG-2) antibody (eBioscience, 14-6504-80, 1:100) and glial fibrillary acidic protein (GFAP) antibody (Invitrogen, PA1-10019, 1:1000), respectively. After primary antibody incubation, samples were washed three times with PBS and then incubated with secondary antibodies and Hoechst 33342 (1:1000, Invitrogen) in B/P solution overnight at 4°C. Afterwards, samples were washed three times with PBS and stored at 4°C until needed.

### Quantification and Statistical Analysis

The analysis for each plot is outlined in the figure legends and/or the corresponding methods mentioned above. In summary, all grouped data are displayed as mean ± standard deviation (SD). All box plots of expression data are represented as median (middle line of the box) ± the 25th percentile (top and bottom lines of the box, respectively). *P* values provided were calculated using one-way ANOVA, followed by Tukey’s t-test. Kaplan-Meier survival curves were generated, and log-rank (Mantel-Cox) analysis was conducted to produce p-values. GraphPad Prism software was employed to create the plots and perform the statistical analysis. Sample sizes for each experiment are specified in the corresponding figures and/or methods above.

### Data and Software Availability

All raw and selected processed data files are available on the NCBI Gene Expression Omnibus (GEO; accession number GSE270481).

## Acknowledgements

This work was supported by National Institutes of Health grants R01NS107462 (to H.Z., R.H.F., and G.D.), R21NS125700 (to H.Z. and R.H.F.), R01NS134159 (to H.Z.), R01NS092735 (to R.H.F.), and NASA 21-3DTMPS_2-0037 (to G.D., H.Z., R.H.F.). Research reported in this publication was also supported by the National Cancer Institute Tisch Cancer Institute Cancer Center Support Grant (P30 CA196521).

---

**Figure S1.**
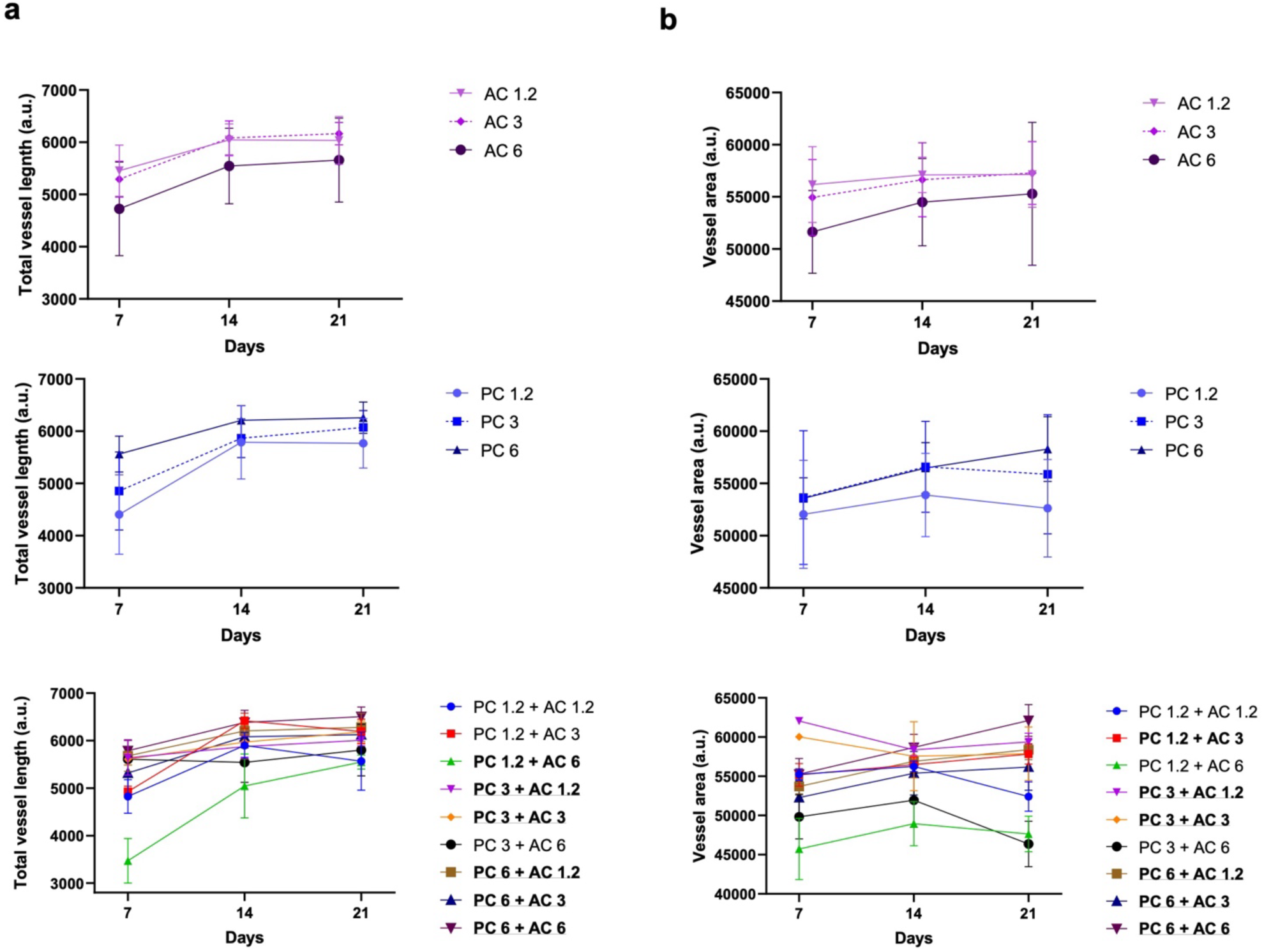
Comparison of astrocyte and pericyte seeding densities to optimize vascular formation in 3D model. (a) Total vessel length in 3D cultures was measured for different seeding concentrations of pericytes (PC) and astrocytes (AC). (b) Total vessel area in 3D cultures was measured for different seeding concentrations of pericytes (PC) and astrocytes (AC). The densities are denoted as 1.2, 3, and 6, indicating 1.2x, 3x, and 6x 10^6^ cells/ml, respectively. Top graphs show seeding of AC or PC alone, respectively, and bottom graphs show combined seeding of AC and PC.

**Figure S2.**
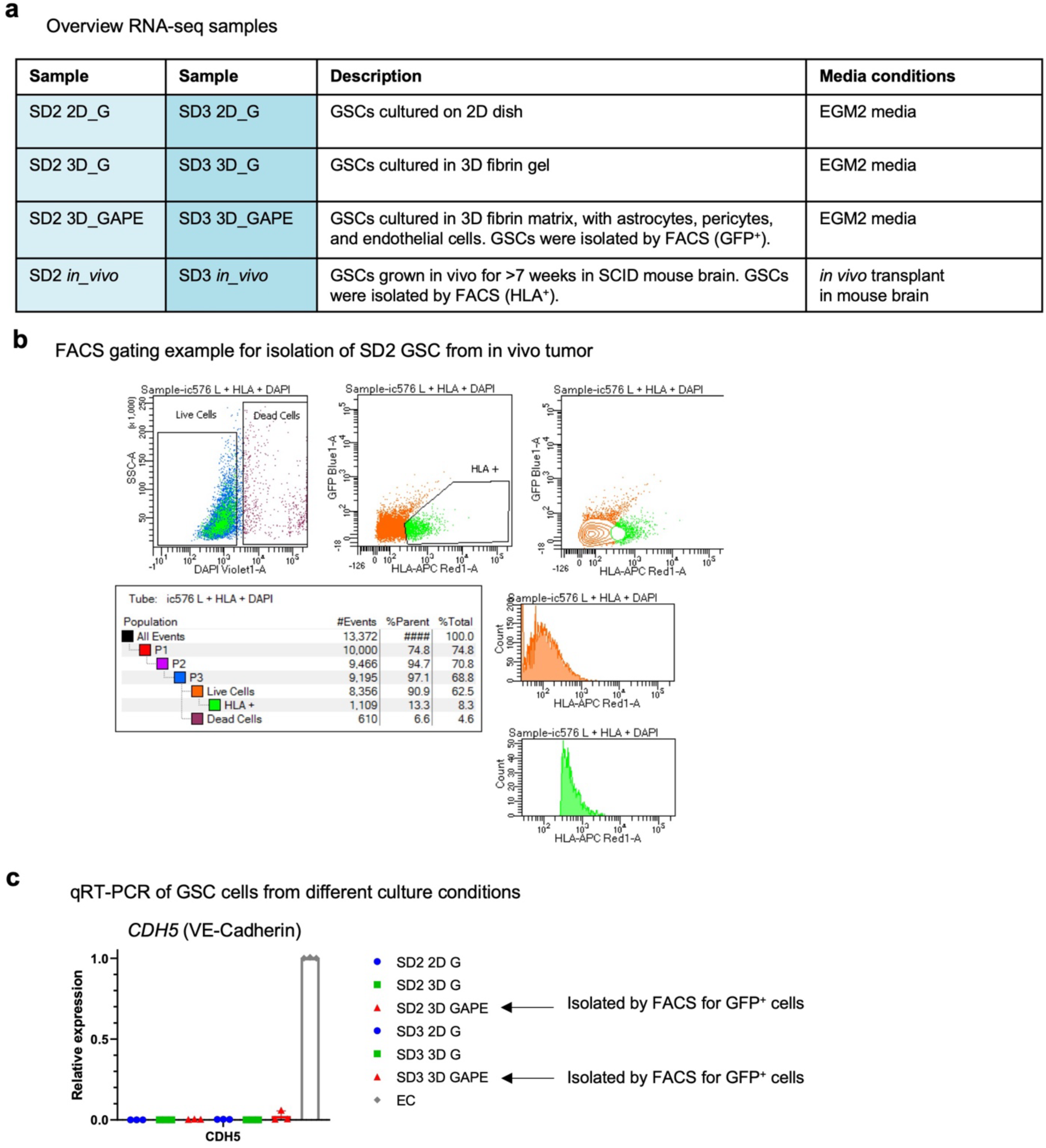
Collection of RNA-seq samples. (a) Overview of sample types and growth conditions. EGM2 media is optimized for growth of endothelial cells. (b) Example of FACS result for isolation of SD2 GSC from a dissociated cell suspension of a brain carrying orthotopic tumor transplant. GBM cells were gated for positive staining with a human-specific anti-HLA antibody (HLA^+^). DAPI dye was used to stain dead cells. (c) qRT-PCR analysis of expression of the endothelial cell (EC) marker gene *CDH5* in GSC isolated from different culture conditions demonstrate purity of FACS isolation from 3D GAPE culture.

**Figure S3.**
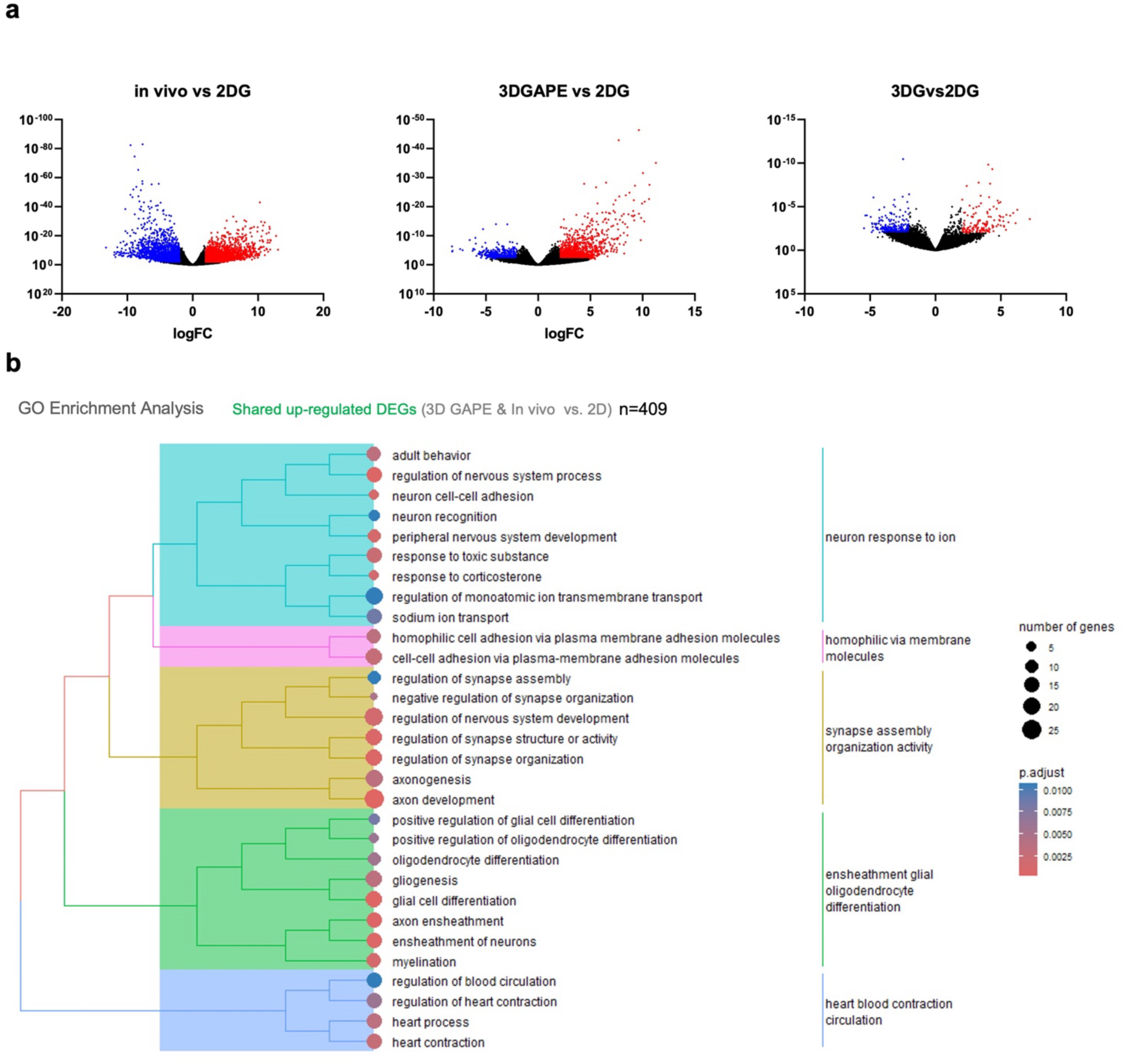
Vulcano plot representations of differentially expressed genes (DEGs). (a) Vulcano plot representations of differentially expressed genes (DEGs) from comparisons of different culture conditions and *in vivo* condition. Cut-offs are *P* < 0.01 and |log2 fold-change| >2. (b) Gene ontology (GO) enrichment analysis of upregulated shared DEGs of *in vivo* and 3D GAPE GSCs vs. 2D.

